# Population genetic structure and connectivity of *Glossina palpalis gambiensis* and *G. morsitans submorsitans* in Southwestern Senegal

**DOI:** 10.64898/2026.09.24.753986

**Authors:** Mame Thierno Bakhoum, Assane Gueye Fall, Adeline Ségard, Momar Talla Seck, Abdou Samath Thiall, Geoffrey Gimonneau, Sophie Ravel, Thierry de Meeûs

**Author notes:** Supervised equally (Sophie Ravel, Thierry de Meeûs).

## Abstract

In sub-Saharan Africa, human and animal health is threatened by African trypanosomoses. These diseases are caused by trypanosomes that are transmitted primarily by tsetse flies. Vector control remains one of the most effective strategies for reducing disease transmission, but requires a minimum level of knowledge of the population biology, ecology and dispersal patterns of the targeted vectors. In this respect, population genetic approaches using appropriate polymorphic genetic markers can provide valuable tools to inform and optimize control strategies. Senegal is currently engaged in a national tsetse control programme and has been identified as a study country within a European Union-funded research initiative, COMBAT (controlling and progressively minimizing the burden of animal trypanosomosis). Baseline entomological studies are currently underway in the Sine Saloum (Southern Senegal). This preliminary study aimed to characterize the population genetic structure and connectivity of tsetse fly species present in Southern Senegal, *Glossina palpalis gambiensis* and *G. morsitans submorsitans*, in order to assess their population sizes (geographically and demographically), population densities and dispersal capacities. The two species exhibited markedly contrasting population structures. The first, *G. p. gambiensis*, displayed an isolation by distance with rather dense subpopulations exchanging immigrants at short distances (1.6 km/generation) and continuously from Sine-Saloum to Casamance, in line with the apparent continuity of the mangrove network found along this coast. In contrast, *G. m. submorsitans* presented a relatively dense population over a 75 km wide area, fragmented into sparsely distributed, very dense and interconnected pockets. These findings indicate that *G. p. gambiensis* likely constitutes a large metapopulation comprising 3266 effective individuals on average (and up to 7765), whereas *G. m. submorsitans* appeared as a single and large population, scattered in small pockets that could be readily replenished by immigrants from distant areas in case of control. These results suggest that elimination is not a realistic option for any of these two populations. Alternatively, localized integrated pest management is likely to be the most effective approach for controlling the negative effects of these biting flies on domestic animals, by reducing trypanosome transmission. However, further entomological investigations and the use of more informative genetic markers are needed to obtain a more precise understanding of the epidemiological processes involved.

## Introduction

Tsetse flies are bloodsucking insects that are restricted to woody environments (riverine or riparian forests, mangroves, dense or protected forests, tree plantations, savannah woodlands and dense thickets) of intertropical Africa (Van den Bossche et al., 2010). They belong to the genus *Glossina* and the superfamily Hippoboscoidea (or Pupiparia) (Krafsur, 2009). This superfamily is characterized by an absence of egg laying, which is replaced by successive larvipositions of a single, full-grown third-instar larva that rapidly pupariate (Krafsur, 2009). For tsetse flies, it occurs approximately every 10 days (Krafsur, 2009). As such, *Glossina* are considered as slowly reproducing animals. They transmit pathogenic trypanosomes to humans and animals. Human African trypanosomosis (HAT), also known as sleeping sickness, is now considered as eliminated as a public health problem (less than 1 case / 10,000 people / year (WHO, 2012), in several countries. It is now targeted by the World Health Organization (WHO) for interruption of transmission (zero human cases for at least five consecutive years (WHO, 2024)) by 2030 (Lejon et al., 2025). Animal African Trypanosomosis (AAT), also known as nagana, continues to represent a heavy burden on sub-Saharan countries with direct and indirect economic losses estimated in billions of dollars each year (Cecchi et al., 2024). Recently, an ambitious programme has been launched by the European Union’s Horizon 2020 research and innovation programme under grant agreement n°101000467, acronym ‘’COMBAT’’ (Controlling and Progressively Minimizing the Burden of Animal Trypanosomosis) (Boulangé et al., 2022). Among the many countries involved in this research programme, Senegal, and particularly its Southern parts, was selected to assess the genetic structure of the two tsetse fly species present in this area. Such information is essential for guiding decisions on the design and implementation of integrated vector control strategies.

Vector control (VC) has been demonstrated as a very useful tool to reduce tsetse populations and tsetse-transmitted trypanosomoses. Local eradications have been successfully achieved through area-wide integrated pest management (AWIPM) using “pour-on” formulations, impregnated targets, and ground- or aerial-based spraying followed by the sterile insect technique (SIT): in the Zanzibar Island (Vreysen et al., 2000) and in the Niayes in Northern Senegal (Vreysen et al., 2021; Seck et al., 2024). The Niayes programme also illustrated the importance of demonstrating sufficient population isolation before implementing an elimination strategy, because sustained immigration from untreated populations can compromise local suppression or eradication efforts (Vreysen et al., 2013). A Chadian project has been recently initiated to eliminate tsetse flies from the sleeping sickness focus of Mandoul using the sterile insect technique (SIT)., with optimistic predictions (Mahamat et al., 2023). In other zones where eradication was not possible, vector control measures, and in particular the use of tiny targets (Rayaisse et al., 2011), can efficiently protect human populations from being bitten by tsetse flies, and as a consequence, against sleeping sickness propagation (Courtin et al., 2015), even when the impact of VC on tsetse populations was shown to be very modest or even negligible (Kagbadouno et al., 2024).

Before the most efficient VC strategy can be determined, the best possible knowledge of the ecology and population biology of the targeted population represents an important prerequisite (Vreysen et al., 2013). To this respect, the study of the spatio-temporal variation of polymorphic genetic markers can prove very efficient, in particular for organisms that are not easy to study directly (De Meeûs et al., 2007). Despite the widespread availability of advanced sequencing technologies, microsatellite markers still represent a cheap alternative and continue to offer a highly accessible and informative approach for resolving population biology parameters of non-model organisms (reproductive strategy, population size and dispersal), and particularly so for arthropod vectors (Prudhomme et al., 2020; Taraveau et al., 2024; Konan et al., 2025; Dorsey et al., 2025).

In this paper, we present a study on the population structure with recently evaluated microsatellite markers (Ravel et al., 2020) of the two species of tsetse flies present in Southern Senegal: *Glossina palpalis gambiensis* and *G. morsitans submorsitans*. Results observed on population densities and dispersal distances per generation are discussed in the context of future vector control programme, and also limitations of the markers used and the need of more performant and easier to use markers such as SSRseq (Lepais et al., 2020) for future studies in other zones.

## Material and methods

### Trap deployment and capture success of tsetse flies

A total of 138 traps were deployed comprising 61 biconical traps (Challier & Laveissière, 1973), 26 Nzi traps (Mihok, 2002), and 51 Vavoua traps (Laveissière & Grébaut, 1990). Each trap was left for three consecutive days and visited daily. Their GPS coordinates, number of *G. p. gambiensis* and *G. m. submorsitans* captured, date of deployment, and apparent densities per trap in each site are indicated in the supplementary Table S1 and Figure S1. Multiple trap designs were used as part of a broader study targeting multiple tsetse species and other hematophagous insects (data not presented in the present paper). Comparisons of capture performances between trap types were undertaken with analyses of variance under R-Commander package (Rcmdr) (Fox, 2005, 2007) of R (R-Core-Team, 2025).

Each tsetse fly individual was sexed and preserved in 70% alcohol in a 1.5 mL tube. In the laboratory, three legs were removed from each of the flies, dried and subjected to chelex treatment in order to obtain DNA for further genotyping.

### Samples of *Glossina palpalis gambiensis* used for the population genetics analyses

The sampling locations are illustrated in the Figure 1. More details are available in the supplementary file S2.

**Figure 1.**
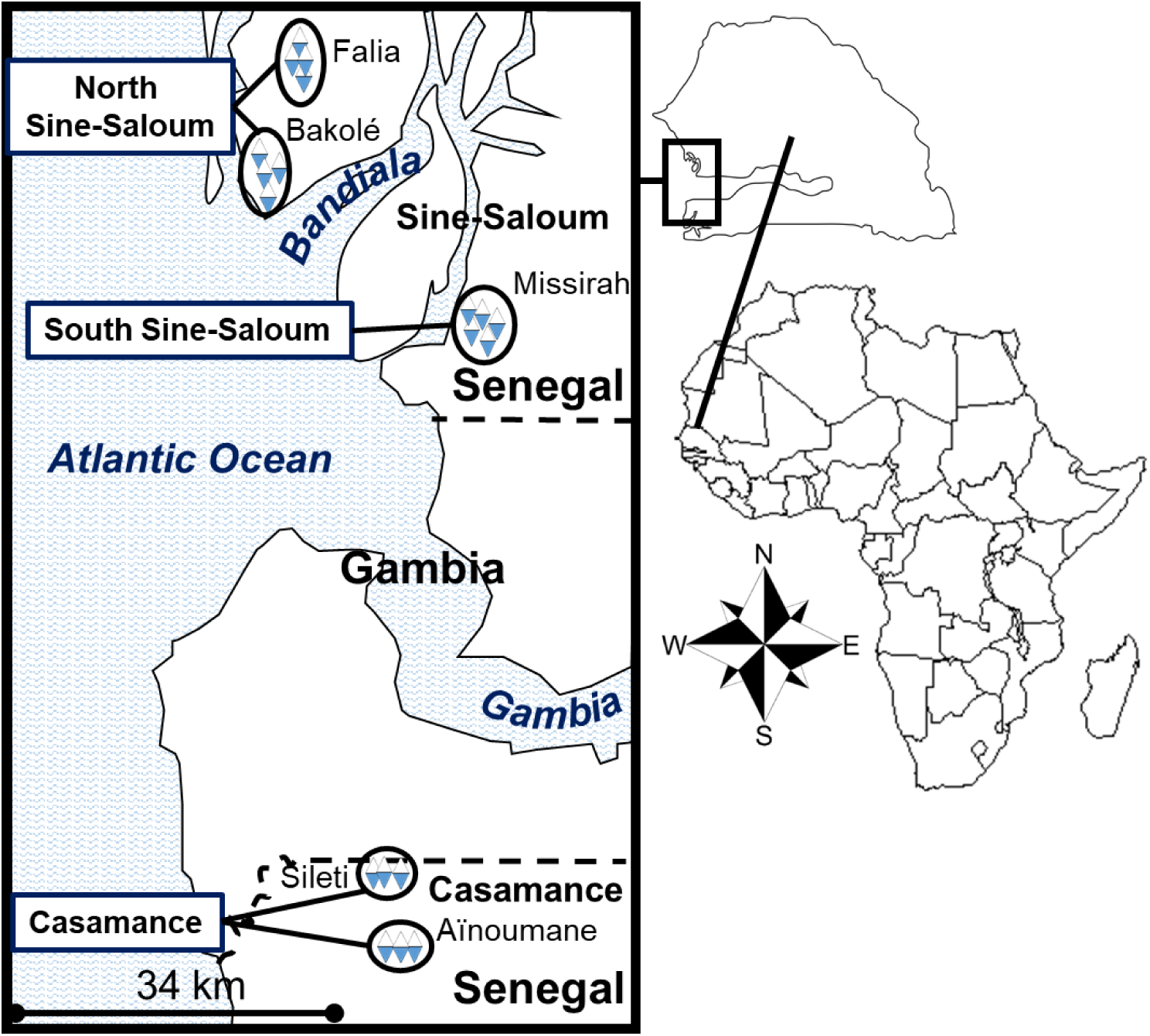
Map of the locations of traps from which *Glossina palpalis gambiensis* specimens were selected for genotyping in Southern Senegal (white and blue diamonds). Sampling sites were named after the nearest village (in black font and in black ellipses). We defined three areas (North Sine-Saloum, South Sine-Saloum and Casamance) (in bold black andsmallest font) and two zones (Sine-Saloum and Casamance in bigger font). Borders between Senegal and Gambia (countries in biggest font) are indicated with black thick dotted lines. Water bodies are named in dark blue (bold and italic font).

### Samples of *Glossina morsitans submorsitans* used for the population genetics analyses

Samples were distributed as in Figure 2.

**Figure 2.**
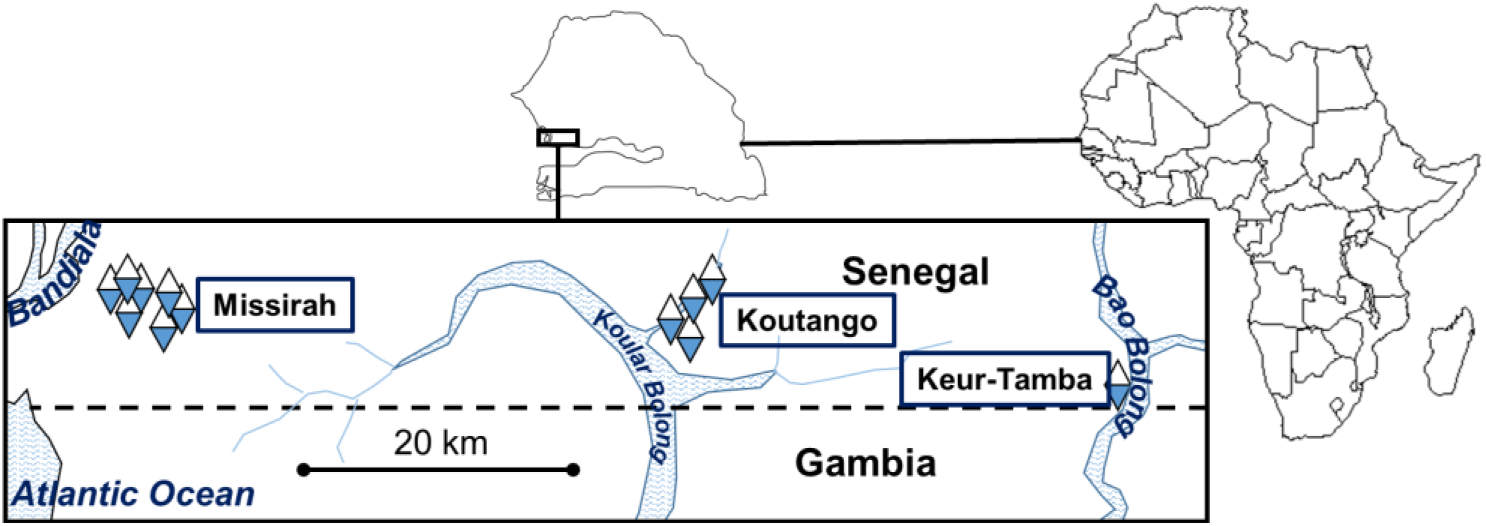
Map of the locations of traps (white and blue diamonds) from which *Glossina morsitans submorsitans* specimens were selected for genotyping, with areas (Missirah, Koutanga and Keur-Tamba) named after the nearest village (squared, in bold font). Border between Senegal and Gambia (countries in biggest font) are indicated with black thick dotted lines. Water bodies are named in dark blue (bold and italic font).

### Genotyping

For *G. p. gambiensis*, we genotyped 147 tsetse flies from 29 traps at nine loci: B3, C102, GPCAG133, PGpg15, PGpg24, PGpg27, XB104, X55.3, and XpGp13 (Ravel et al., 2020), three of which were X-linked and thus haploid in males (indicated with the X as 1^st^ letter). We first studied only the 6 autosomal loci for all individuals (dataset called autosomes hereafter). We also checked if our results matched those obtained with X-linked loci in females only (dataset called females hereafter). All flies were collected between the fall of December 2022 and February 2023. According to the theoretical life cycle of tsetse flies (Artzrouni & Gouteux, 2006), pupal stage lasts from 20 to 100 days, the pre-reproductive adult stage 20 days, and the time between two larvipositions is 10 days. The average generation time should thus be between 2 and 5 months. We then assumed that the individuals sampled during this survey belonged to the same cohort. Genotypes are available in the Supplementary File S2.

For *G. m. submorsitans*, we genotyped 190 flies from 12 traps at only six loci: Gmm20291, GmmK06, Gmm26134, GmsCAG29B, GmmH09, and GmmA06 (Ravel et al., 2020). Data are available in the supplementary file S3.

Genotyping at each locus and for each species was obtained following the detailed protocols published by Ravel et al. (2020).

### Data analyses

#### Finding the most relevant level of subdivision

To describe the structure of the populations we used Wright’s *F*-statistics (Wright, 1965). The deviation from local random mating between individuals is described by the inbreeding of individuals relative to the inbreeding of subpopulations (or more exactly subsamples) (*F*_IS_). It varies from negative values (excess of heterozygotes) down to −1, 0 (panmixia) to positive values (homozygote excesses) up to 1. Subdivision was assessed with the inbreeding of subsamples as compared to total inbreeding (*F*_ST_), which varies from 0 (no subdivision) to 1 (all subsamples fixed for one or the other allele, i.e. complete isolation between subsamples). Finally, inbreeding of individuals as compared to total inbreeding (*F*_IT_) is produced by the combined effect of the two first: *F*_IT_=1-(1-*F*_IS_)(1-*F*_ST_). These parameters were estimated with Weir and Cockerham unbiased estimators (Weir & Cockerham, 1984): *f* (*F*_IS_), *θ* (*F*_ST_) and *F* (*F*_IT_). Note that the estimator of *F*_ST_, unlike the parameter, is centered on 0 under the null hypothesis (no subdivision), and can thus display negative values. Strongly negative estimates of *F*_ST_ would then correspond to subsamples that are genetically more similar than if there would have been randomly sampled from the same subpopulation. We also computed 95% confidence intervals of these statistics (95%CI) with 5,000 bootstraps over loci. To test for the significant deviation from 0 of *F*_IS_, we randomized alleles between individuals, within subsamples and compared randomized *f*’s to the observed one. For subdivision, we randomized individuals between subsamples, and used the *G*-based test (Goudet et al., 1996) for each locus and over all loci. We also tested the significance of linkage disequilibrium (LD) with the *G*-based randomization test combined over subsamples. Combining the *G*’s over loci or subsamples represents the most powerful method (De Meeûs et al., 2009). For all these procedures, we implemented 10,000 randomizations. These estimates and testing were undertaken with Fstat 2.9.4 (Goudet, 2003), updated from Fstat 1.2 (Goudet, 1995). The default test for panmixia being one-sided (*p*_1_, *F*_IS_>0), we undertook a two-sided conversion by doubling the appropriate probability: *p*_2_=2×*p*_1_ for *p*_1_>0.5, and *p*_2_=2×(1-*p*_1_) otherwise.

To define the level at which significant subdivision occurs, we used Goudet et al’s 1994 *F*_IS_ based method (Goudet et al., 1994), extended to *F*_ST_ and to the proportion of significant locus pairs in significant LD (%LD).

For *G. p. gambiensis*, we defined five hierarchical levels (see Figure 1): the trap (T); the site (S) (Falia, Bakolé, Missirah, Sileti and Aïnoumane); the area (A) (North Sine-Saloum, South Sine-Saloum and Casamance); the zone (Z) defined by the Northern part (Sine-Saloum) and the Southern part (Casamance) of Senegal, and the total (not studied thoroughly).

For Gms, we defined only three levels: Trap, Area and the Total (AllInOne) (see Figure 2).

Four (at least) corresponding *F*_IS_, *F*_ST_ and %LD could be compared: *F*_IS-T_, within traps; *F*_IS-S_, within sites (ignoring traps); *F*_IS-A_ (pooling all individuals from the area into one subsample); *F*_IS-Z_ (for *G. p. gambiensis*, pooling all individuals from Sine-Saloum on one side, and those from Casamance on the other side), and, for *G. m. submorsitans*, *F*_IS-AllInOne_ (all individuals pooled into a single sample). If individuals that do not belong to the same demographic unit are pooled, this should increase *F*_IS_. This was tested with a one-sided Wilcoxon signed rank test for paired data (paired by loci) with the alternative hypotheses: *F*_IS-T_<*F*_IS-S_<*F*_IS-A_<*F*_IS-B_<*F*_IS-Z_<*F*_IS-AllInOne_. These tests were done with Rcmdr. We also compared *F*_ST_ with the same method but with the reverse order of alternative hypotheses, as pooling individuals from genetically distant population tends to dilute the effect of subdivision (De Meeûs, 2018). We compared the proportion of significant LD tests (%LD) between sampling strategies with one sided Fisher exact tests, as %LD is expected to increase with a moderate Wahlund effect (Manangwa et al., 2019) (command fisher.test in R). For a last confirmation of the most relevant hierarchical level, we also undertook tests of subdivision between paired subsamples at the required level. All paired tests produced series of tests with some degree of dependency. This would suggest using Benjamini and Yekutieli (BY) (Benjamini & Yekutieli, 2001) adjustment. Contrarily to what we usually undertook in previous population genetics articles undertaken by one of us (TdM), and for reasons expounded in Appendix A, we used Bejamini and Hochberg’s (BH) correction (Benjamini & Hochberg, 1995) instead. This procedure is easily undertaken with the R command "p.adjust(c(*p*_1_, *p*_2_,… *p*_k_), method="BH")", where c(*p*_1_, *p*_2_,… *p*_k_) is the vector containing the *k p*-values to combine.

#### Quality tests of loci

In case of heterozygote deficits (*F*_IS_>0), we undertook several techniques to detect amplification problems (De Meeûs, 2018; Manangwa et al., 2019; De Meeûs & Noûs, 2022).

Null allele frequencies were assessed with the EM algorithm (Dempster et al., 1977) with FreeNA (Chapuis & Estoup, 2007). We computed the average frequencies of null alleles across subsamples for each locus, using subsample sizes as weights. More accuracy can be obtained when missing data are recoded as homozygotes for null alleles’ (i.e. 999999), if missing data indeed correspond to null homozygotes. Some loci may display a number of missing data (*N*_miss_) that appears bigger as compared to the *F*_IS_ they display. In such case, we left missing data as such (i.e. coded "0"). This was checked first by the regression *F*_IS_∼*N*_miss_. Typically, loci located far below the regression line display too much missing genotypes, which probably do not correspond to null homozygotes. For instance, missing data at loci with a negative *F*_IS_ can hardly correspond to null homozygotes. With null allele frequency estimates (*p*_nulls_), we computed the number of expected missing data as E(*N*_nulls-*i*_)=*p*_nulls-*i*_²×*N_i_*, where *i* stands for subsample *i*. For each locus, we compared the sum of these expected values across subsamples and compared to the observed number of missing genotypes with a one-sided (less) exact binomial test with R (command binom.test), for each locus, and corrected with the BH procedure with R (command p.adjust). We quantified the goodness of fit of the model under panmixia with null alleles with the regression *F*_IS_∼*p*_nulls_. We interpreted the determination coefficient (*R*²) as the proportion of *F*_IS_ variation explained by null alleles. We considered a value *R*²>0.9 as reasonably satisfactory, and that *F*_IS_ was entirely explained by null alleles when *R*²>0.95. Other values required more investigations and/or discussions. We undertook these regressions with a spreadsheet programme. The intercept of the regression (*F*_IS-0_) provided the probable value of this statistic in absence of null alleles.

Stuttering detection was undertaken with De Meeûs and Noûs method (De Meeûs & Noûs, 2022). For short allele dominance (SAD) detection, we tested the negative correlation between *F*_IT_ and allele sizes (*A*) (Manangwa et al., 2019). In case of doubt, and to avoid the distortion effect of rare alleles, we also undertook the regression *F*_IS_∼*A* weighted by *p_a_*×(1-*p_a_*), and where *p*_a_ is the frequency of allele *a* overall subsamples (De Meeûs et al., 2004).

To compute 95%CI of *F*_IS_ for each locus, we also undertook 5,000 bootstraps over individuals with Genetix (Belkhir et al., 2004). The average across subsamples were computed for each locus as:

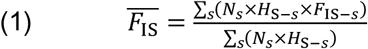

where *N_s_*, *H*_S-*s*_ and *F*_IS-*s*_ are the number of visible genotypes in subsample *s* for a given locus, Nei’s unbiased estimate of genetic diversity and Weir and Cockerham unbiased estimator of *F*_IS_ in that subsample, as computed by Genetix as well.

We also checked that all loci used were statistically independent with the *G*-based randomization test described above between each locus pair. For *L* loci, this thus represents a series of *k*=*L*(*L*-1)/2 tests where we needed to discriminate which *p*-values are significant by chance. There is some degree of dependency in such series, but for the same reasons explained above, we also used BH to adjust LD *p*-values.

When relevant, we estimated the rate of systematic mating between full sibs as *s*_sib_=4*F*_IS_/(1+3*F*_IS_) (Chevillon et al., 2007).

#### Genetic subdivision

As mentioned above, subdivision was first assessed over all subsamples with Weir and Cockerham’s unbiased estimator of *F*_ST_. In case of null alleles, we corrected this estimate with the ENA algorithm with FreeNA (*F*_ST-ENA_) with its 95%CI (5,000 bootstraps over loci). However, due to the excess of polymorphism of microsatellite markers, *F*_ST_ can be underestimated. This occurs when the correlation between *F*_ST_ and *H*_S_ is significantly negative (Wang, 2015), and where *H*_S_ is the unbiased estimator of local genetic diversity (Nei & Chesser, 1983). In that case only, we corrected our estimates with Meirmans’ method and its software RecodeData (Meirmans, 2006) to obtain the maximum possible *F*_ST_ with our data (*F*_ST-max_). This quantity was then used to obtain a standardized estimate *F*_ST-ENA_’=*F*_ST-ENA_/*F*_ST-max_. We computed the number of immigrants as *N_e_m*=(1-*F*_ST_)/(4*F*_ST_) with the relevant corrected version of *F*_ST_ (Ravel et al., 2023).

We also assessed isolation by distance with Rousset’s regression model in two dimensions, *F*_R_=*a*+*b*×ln(*D*_geo_), where *F*_R_=*F*_ST/_(1-*F*_ST_) (using *F*_ST-ENA_ in case of null alleles), *a* is the intercept, and *b* the slope of the regression and *D*_geo_ is the geographic distance between two subsamples (Rousset, 1997). If the regression is significant, the reverse of the slope provides the neighborhood of a subpopulation (roughly the number of connected individuals): 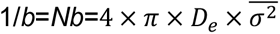. In this equation, *D_e_* is the effective population density of the population, i.e. the number of adults that can transmit their genes divided by the surface occupied (*D_e_*=*N_e_*/*S*), and 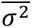 is the average of the squared axial distance between adults and their parents (the axial distance is half the distance between adults and their parents, *δ*). If *D*_e_ is known, and if the distribution of axial distances is not too skewed and/or kurtosis not too pronounced, this can also lead to estimate the dispersal distance per generation (Séré et al., 2017):

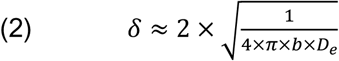

In case of excessive skewness/kurtosis, an overestimate of twice or thrice the real value can be expected, but the order of magnitude cannot really change.

Rousset has also shown that, in two dimensions, the number of immigrants exchanges by neighbors was *N_e_m*=1/(2×*π*×*b*).

We admitted a significant isolation by distance signature when all three slopes of the model (average, and the two 95%CIs of bootstraps) were positive. When one of the 95%CI slopes appeared moderately negative, we checked the signature of a weak (if any) isolation by distance with a Mantel test (Mantel, 1967) between the two matrices defined by the geographic distances between paired subsamples and the chord distance of Cavalli-Sforza and Ewards (Cavalli-Sforza & Edwards, 1967), computed with the INA correction for null alleles (*D*_CSE-INA_) with FreeNA. This method may indeed prove more powerful in some instances (Séré et al., 2017). To undertake the Mantel test, we used Fstat, which computes Pearson’s correlation (*r*) between the two matrices, and with 10,000 randomizations. The output of Fstat is two-sided. We thus converted the *p*-value into a one-sided *p*-value for a positive correlation by halving it (in case of positive correlation of course).

The geographic distance between each subsample pairs (*D*_geo_) was computed after the GPS coordinates in decimal degrees with the package geosphere (Hijmans et al., 2019) of R (see Appendix B).

#### Effective population sizes

We used five methods. The heterozygote excess method (Hex) (De Meeûs & Noûs, 2023), the linkage disequilibrium method (LD) (Waples, 2006) corrected for missing data (Peel et al., 2013), the co-ancestry method (CoA) (Nomura, 2008), the one and two loci correlation (1L2L) (Vitalis & Couvet, 2001a) and the sibship method (Sib) (Wang, 2009). For Hex we computed *N_e_* for each locus and each subsample, and computed the average across loci, in each subsample (De Meeûs & Noûs, 2023). For LD and CoA we used NeEstimator (Do et al., 2014). For 1L2L we used Estim (Vitalis, 2002), updated from Vitalis and Couvet (2001b). Finally, Sib *N_e_* were computed with Colony (Jones & Wang, 2010).

We computed the average across subsamples for each method, with the minimum and maximum, values (minimax) and the number of usable values (uv) (other than "Infinite"). The grand average was then computed across methods, with uvs as weights (De Meeûs & Noûs, 2023).

#### Dispersal and effective population densities

We used the number of immigrants and effective population size computed above to estimate the proportion of immigrants exchanged between each of our subsamples as *m*=*N_e_m*/*N_e_*. We used the average GPS coordinates across traps within the same subsample unit used to compute geographic distances between each subsample pairs (*D*_geo_) and obtained the dispersal distance per generation for each subsample pair as *δ*=*m*×*D*_geo_. As the surface occupied by a population of *G. p. gambiensis* was uneasy to define, we used the equation (2) to compute *D_e_*=1/(*δ*²×*π*×*b*) (only in case of significant isolation by distance). We then could define the average surface occupied by a subpopulation as *S*=*N_e_*/*D_e_*. Finally, we also defined one other surface, *S*_ETT_, defined by the four most extreme Eastern, Southern, Western and Eastern traps with at least one fly. This surface was computed with geosphere. We then used *S*_ETT_ to compute the total effective population size of tsetse flies contained in the investigated polygon as *N_e_*_-tot_= *S*_ETT_×*D_e_*. This figure helped us to get a rough perception of the total number of subpopulations contained in the zone we investigated as *n*=*N_e_*_-tot_/*N_e_*. Please, note that negative *F*_ST_’s were interpreted as an absence of subdivision and translated into *m*=1.

For *G. m. submorsitans*, two traps (MB_P1 and KTM_P1) (see Supplementary File S1), were both most Western and Northern, for the first, and most Eastern and Southern, for the second. We defined *S*_ETT_ as the rectangle, the diagonal of which was defined by these two traps. We also computed the surface *S*_external_ of the polygon defined by most external traps of each area: MBP1, MBP_4, PB_P3, FB_P3, FB_P1, and RF_P5, in Missirah; FST_P9 and RS_P9, in Koutangan; and KTM_P1 in Keur-Tamba (see Supplementary File S1). These surfaces were computed with geosphere (see Appendix B).

We also tested for a genetic signature of sex-biased dispersal with the R package HierFstat (Goudet, 2005) (see scripts in Appendix C). For this, we used the four available statistics: the corrected assignment index and its variance (*AI*_c_ and *vAI*_c_) (Favre et al., 1997), and the unbiased estimators of wright’s *F*_ST_ and *F*_IS_. The most philopatric gender should display the highest *AI*_c_, the smallest *vAI*_c_, the highest *F*_ST_ and the smallest *F*_IS_. Significance was assessed through randomizations of gender within each subsample, keeping the sex-ratio unchanged. The number of randomizations was 10,000 for *AI*_c_ and *vAI*_c_ and only 1,000 for *F*_ST_ and *F*_IS_ as the procedure took too much time with those parameters.

## Results

### Trap performances

For both species, trap types did not significantly influence the success of capture of flies in any of the site (*p*-values>0.2) (Table 1). On the other hand, sites displayed strong differences (Tables 1 and 2).

**Table 1.**
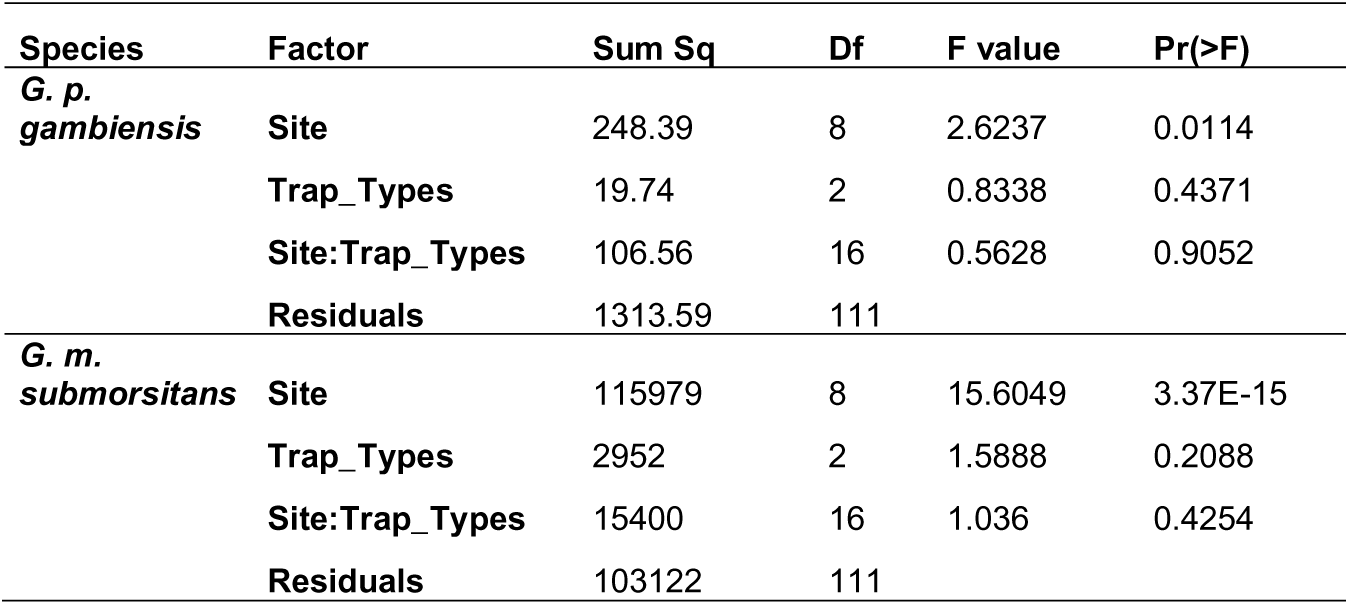
Results of the analyses of variance for the abundances of *Glossina palpalis gambiensis* (*G. p. gambiensis*) and *G. morsitans submorsitans* (*G. m. submorsitans*) in different traps and sites of Southwestern Senegal.

**Table 2.** Averages (mean), standard deviations (sd) of the number of flies captured by different traps (*N*_traps_) for *Glossina palpalis gambiensis* (*G. p. gambiensis*) and *G. morsitans submorsitans* (*G. m. submorsitans*) in Southwestern Senegal.

| Site | *G. p. gambiensis* | | *G. m. submorsitans* | | $N_{traps}$ |
| --- | --- | --- | --- | --- | --- |
|  | mean | sd | mean | sd |  |
| Ainoumane | 0.5 | 1.0 | 0 | 0 | 10 |
| Bakole | 3.3 | 7.3 | 0 | 0 | 19 |
| Falia | 1.2 | 1.5 | 0 | 0 | 10 |
| Keur_Nala | 0 | 0 | 0 | 0 | 17 |
| Keur_Tamba | 0 | 0 | 0.9 | 2.5 | 8 |
| Koutango | 0 | 0 | 3.2 | 4.6 | 20 |
| Missirah | 3 | 4.4 | 85 | 82 | 19 |
| Ndiamacouta | 0 | 0 | 0 | 0 | 25 |
| Sileti | 2.3 | 3.2 | 0 | 0 | 10 |

### Relevant hierarchical level of subdivision

For *G. p. gambiensis*, with the six autosomal loci, there was a significant subdivision (*p*-value<0.0001), whatever the sampling design. According to Figure 3, no comparison stayed significant after BH correction, except with *F*_IS_ for which significant subdivision began between sites. As a confirmation, subdivision test between paired sites confirmed this interpretation. Indeed, all tests were significant after BH correction (all *p*_BH_≤0.0022). Consequently, we kept sites as the relevant subdivision level for subsequent analyses.

**Figure 3.**
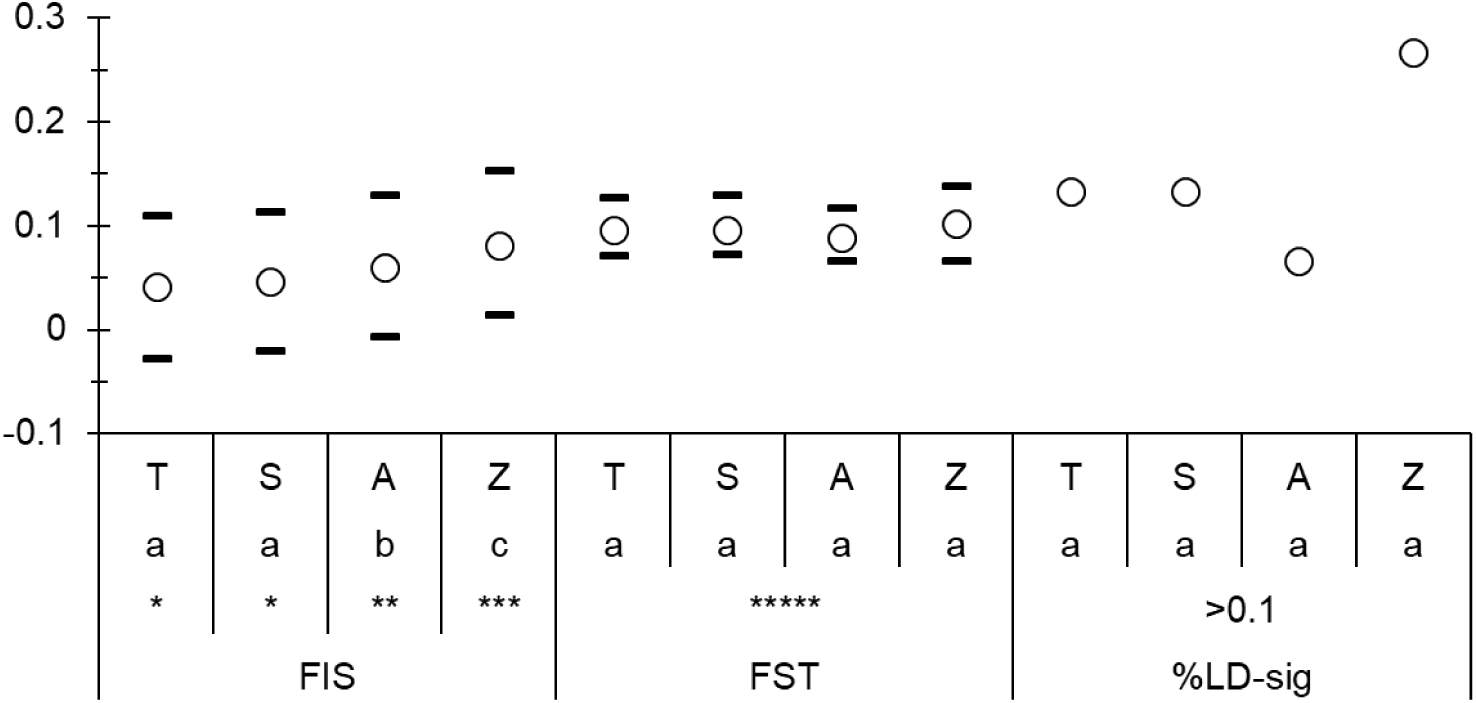
Test comparisons of *F*_IS_, *F*_ST_ and %LD for different sampling designs: T (Traps), S (Sites), A (Areas), and Z (Zones) as defined in the text and in Figure 1, for *Glossina palpalis gambiensis* from Sine-Saloum and Casamance (Southern Senegal). Sampling designs that differ at the 5% significance (i.e. *p*-value<0.05) are indicated with different letters (a, b or c) (after BH correction). The level of significance of the parameters are also indicated: * (*p*-value<0.05), ** (*p*-value<0.01), *** (*p*-value<0.001), and **** (*p*-value<0.0001). Data were with 6 autosomal loci.

For *G. m. submorsitans*, no test appeared significant, even for the *F*_IS_ when all individuals of all areas were pooled into a single unit (i.e. *F*_IS-traps_=*F*_IS-AllInOne_=0.353). It would appear that the real level for a subpopulation of *G. m. submorsitans* would be the whole zone sampled (at least). Nevertheless, and unless specified otherwise, we kept the level area with three subpopulations: Missirah, Koutanga and Keur-Tamba to undertake most subsequent analyses.

### Quality tests of loci

There was a modest though significant and variable, heterozygote deficit: *F*_IS_=0.047 in 95%CI=[-0.021, 0.113] (Two-sided *p*-value=0.0228) in sites for all individuals from *G. p. gambiensis* using autosomal loci (Figure 4). This deficit did not stay significant in females (all loci) (*F*_IS_=0.032 in 95%CI=[-0.024, 0.091], *p*-value=0.1666).

**Figure 4.**
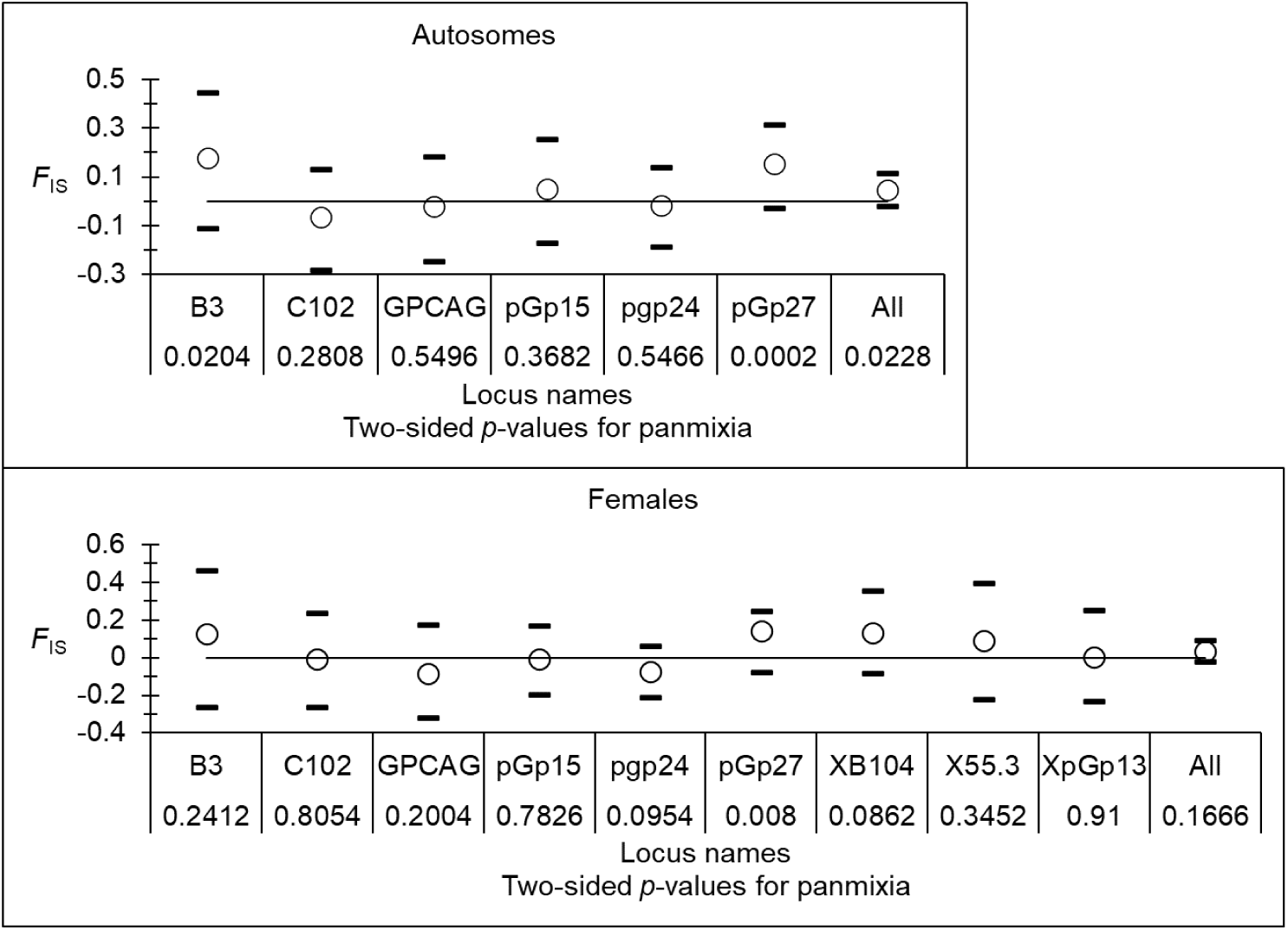
Unbiased estimates of Wright’s *F*_IS_ for each locus and overall (All), and, with their 95% confidence intervals, for *Glossina palpalis gambiensis* from Sine-Saloum and Casamance (Senegal). Results of significant deviations above or under 0 (two-sided *p*-values for panmixia) are also indicated. Results obtained for all individuals with autosomal loci only are given on the top, while results observed at all loci in females only are in the bottom of the figure.

For *G. m. submorsitans*, there was a more important, highly variable and highly significant heterozygote deficit within areas: *F*_IS_=0.353 in 95% CI=[0.134, 0.571] (*p*-value<0.0002) (Figure 5).

**Figure 5.**
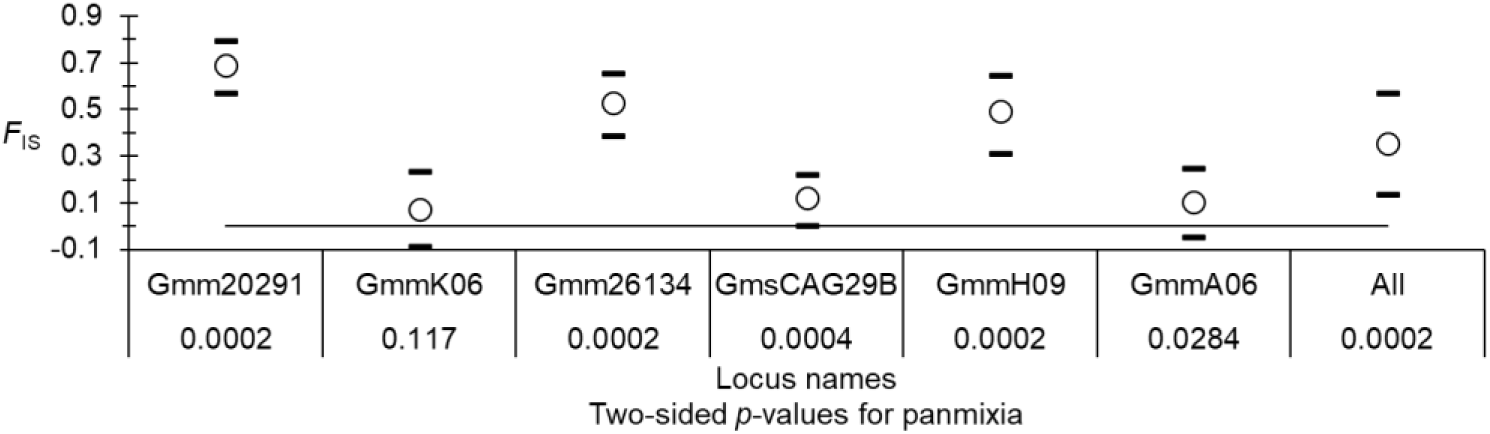
Unbiased estimates of Wright’s *F*_IS_ for the six loci and over all, with their 95% confidence intervals, for *Glossina morsitans submorsitans* from the three areas, Missirah, Koutanga and Keur-Tamba (Senegal). Results of significant deviations above or under 0 (two-sided *p*-values for panmixia) are also indicated.

For *G. p. gambiensis*, after several attempts, it appeared that the best coding for missing data was null homozygotes (999999) for B3, pGp27, XB104, X55.3 and XpGp13, and missing genotypes (0) for C102, GPCAG, pGp15 and pGp24. We found that null alleles explained well the heterozygote deficit and its variation across loci. The regression *F*_IS_∼*p*_nulls_ provided rather good results (*R*²=0.8907, *F*_IS-0_=-0.0315 in 95%CI=[-0.2347, 0.1547] for autosomes; *R*²=0.8813, *F*_IS-0_=-0.0498 in 95%CI=[-0.2591, 0.1587] for females). Three loci presented a lot of missing genotypes many of which probably did not correspond to null homozygotes (C102, pGp15 and XB104). These observations are compatible with pangamic subpopulations with a rather small effective population size *N_e_*=10-16 in 95%CI=[2, Infinite] (*F*_IS_ based *N_e_*) (De Meeûs & Noûs, 2023).

For *G. m. submorsitans*, the best coding of missing data was 999999 for loci Gmm20291, Gmm26134 and GmmH09 and missing genotypes (0) for Gmm06, GmsCAG29B and GmmA06. Null alleles explained 95.5% of the variance of *F*_IS_ across loci, but with a positive intercept (*F*_IS-0_=0.0567 in 95%CI=[-0.0925, 0.1957]). Moreover, one locus, Gmm26134, displayed a highly significant stuttering signature (*p*-value=0.0015). Due to the continuity of allele sizes at this locus, and despite our attempts with arbitrary cutoffs, pooling alleles to cure this locus for stuttering, as described elsewhere (De Meeûs et al., 2021; De Meeûs & Noûs, 2022), revealed impossible. We thus excluded this locus. With the five remaining loci, the regression *F*_IS_∼*p*_nulls_ displayed a *R*²=0.9914, with still a positive intercept: *F*_IS-0_=0.0454 in 95%CI=[-0.1034, 0.1852]. This positive intercept, which seemed not significantly different from 0, could hardly come from a Wahlund effect, given our results within traps. It may then come either from a very big effective population size, with an important variance estimate due to modest sample size and number of loci, and/or to systematic mating between sibs, with a rate *s*_sib_≈0.2 in 95%CI≈[0, 0.5].

### Subdivision

For both species, Wang’s criterion was negative (no significant effect of genetic diversity on *F*_ST-ENA_). For *G. p. gambiensis*, combining results for the 6 autosomes with those of females on X-linked markers, Spearman’s correlation was *ρ*=-0.4333 (*p*-value=0.125), and for *G. m. submorsitans*, *ρ*=-0.3 (*p*-value=0.3417) (for the remaining five loci). We thus did not use correction for polymorphism to estimate subdivision measures.

According to the slopes of the regression between *F*_ST-ENA_ based Rousset’s genetic distance and the natural logarithm of genetic distances (ln(*D*_geo_)), we observed a marginally not significant isolation by distance between sites of *G. p. gambiensis*: slope *b*=0.026 in 95%CI=[-0.0009, 0.0281] with autosomes, but significant with the Mantel test (*p*-value=0.0145), and slope *b*=0.02 in 95%CI=[-0.0026, 0.0273] with females, but also significant with the Mantel test (*p*-value=0.0386). This translated into a neighborhood size of *Nb*=50 in 95%CI=[37, infinite] connected individuals and *N_e_m*=8 in 95%CI=[6, undefined] for females and *Nb*=38 in 95%CI=[36, infinite] and *N_e_m*=3 in 95%CI=[3, undefined] with autosomes.

For *G. m. submorsitans*, subdivision was globally not significant (*p*-value=0.223), with a small *F*_ST-ENA_=0.005 in 95%CI=[-0.0013, 0.0126]. We thus assumed that tsetse flies diperse freely across the whole zone (i.e. around 80 km long), during a life time (i.e. two to five months).

### Effective population sizes

In *G. p. gambiensis* populations, effective population sizes ranged between *N_e_*=64 in minimax=[9, 170] considering all individuals and autosomal loci, and *N_e_*=19 in minimax=[5, 35] in females and all loci. This difference probably comes from the fact that heterosomal markers are haploid in males and thus drift faster than autosomal ones. We thus kept data from autosomal markers for dispersal and density inferences. Please note that, with autosomes, the effective population size was in the same order of magnitude as neighborhood size computed above.

For *G. m. submorsitans*, we chose to present results obtained when considering all individuals within a single population. It provided a rather large effective population size averaged across methods: *N_e_*=724 in minimax=[27, 1425].

For both species, several "infinite" values could not be used to compute the weighted averaged *N_e_*, depending on the method and the site. This means that the figures obtained probably represent underestimates. The strategy proposed by Waples (2024), using harmonic means, and setting "infinite" results at (say) 10,000, did not help here as the results provided even smaller average effective population sizes as the one obtained with the arithmetic mean (and ignoring "infinire" results). We thus kept the "underestimated" values obtained with weighted arithmetic means as described by De Meeûs and Noûs (2023).

### Dispersal and effective population densities

For *G. p. gambiensis*, we used the *F*_ST-ENA_ and 95 %CI between each pair of sites for autosome data to compute *N_e_m* and use *N_e_* and its minimax to extract the average dispersal distance *δ*=1.6 km/generation in 95%CI=[1, 3.2], and in minimax=[0.5, 21] km/generation. This led to rather important effective population densities: *D_e_*=5 tsetse flies/km² in 95%CI=[1, 13], and in minimax=[0.03, 90.5] flies/km² (ignoring infinite results). This led to an average surface of *S*_site_=12 km² in 95%CI=[5, 48], and in minimax=[2, 328] km² (ignoring infinite results). For the record, if we use the maximum distance between two traps that captured a fly in each site as a measure of the diameter of a sampled site, this gives an average surface of *S*_site-sampled_=2.5 km², in minimax=[0.03, 5.3]. Most extreme traps of the whole sample defined a surface of *S*_ETT_=605 km² for *G. p. gambiensis* captured during this survey. This suggested a total effective population size of *N_e_*_-tot_=3266 in 95%CI=[813, 7765] individuals, and ranging in minimax=[16, 54736] flies. This would also mean that an average of 51 subpopulations of *G. p. gambiensis* in 95%CI=[13, 122] are distributed in the investigated region.

For *G. m. submorsitans*, the two surfaces we defined were *S*_ETT_=705 km² and *S*_external_=89 km². This allowed computing that the global effective population densities were relatively small, with *D_e_*_-ETT_=1.03 flies/km² in minimax=[0.04, 2.02], or *D_e_*_-external_=8.2 flies/km² in minimax=[0.3, 16].

For gender specific genetic structure, in *G. p. gambiensis*, all parameters pointed in the same direction (male biased dispersal), but none significantly so (all *p*-values>0.18).

Given the absence of subdivision signature in *G. m. submorsitans*, we considered that looking for a sex-biased dispersal signature was irrelevant in that species at the geographic scale investigated.

## Discussion

This study aimed to characterize the population genetic structure and connectivity of tsetse fly species present in southern Senegal, with the goal of informing vector control strategies in an area targeted by national and international intervention programmes. Our results reveal strikingly contrasting population structures between *Glossina palpalis gambiensis* and *G. morsitans submorsitans*.

We evidenced a rather viscous environment for *G. p. gambiensis*, with a significant isolation by distance with rather short dispersal distances of 1 to 3 km/generation (computed with averaged *N_e_*) average, and in a maximum range between 500 m and 21 km. Effective population sizes matched expectations made from neighborhood sizes and suggested the existence of multiple interconnected subpopulations dispersed in the whole zone concerned, from the Sine-Saloum to Casamance. The meta-population defined by these subpopulations includes those that are known to exist in Gambia (De Meeûs et al., 2015), which remained unexplored in the present paper. Indeed, an absence of subpopulations between Missirah and Casamance would have produced a brutal jump of genetic differentiation. This would have led to a visible break in the isolation by distance slope that we did not observe during our analyses. Effective population densities appeared relatively important, leading to the probable existence of a large total effective population size across the whole zone roughly ranging between 1,000 flies and 8,000 flies, exchanging a few flies per generation between direct neighbors. Nevertheless, many other subpopulations probably exist around the investigated surface defined by the traps we deployed, and this figure is probably a rather important underestimate of the true total effective population size of *G. p. gambiensis* in Southern Senegal.

Compared to other studies, the dispersal distances observed for *G. p. gambiensis* are close to those found for the same species but in a different environment in Burkina-Faso with similar markers (Bouyer et al., 2009; Kone et al., 2011) or with the Mark-Release-Recapture technique (MRR) (Bouyer et al., 2009), with very variable densities (0.2 and 200 flies per km²). Nevertheless, it appeared very different from what was observed in a similar environment in the Mangrove of Guinea, where dispersal distances of at least 40-50 km per generation was inferred by similar markers (Kagbadouno et al., 2024) and SSR-Seq loci (De Meeûs et al., 2025), with much smaller densities (0.5 flies /km² at best). Nevertheless, the mangrove population studied by Kagbadouno et al. is known to be strongly disturbed by vector monitoring and control since 2011. This may have triggered large distance dispersal (De Meeûs et al., 2019). Comparison with more ancient data is not easy. In 2005 (Solano et al., 2009), the genetic differentiation between Dubreka (mangrove) (*N_e_*=1000) and the remote savannah site Falessadé (46 km away inland Guinea) (*N_e_*=40) suggested a much smaller dispersal (*δ* between 1 and 4 km/generation). Comparision is even more difficult as Falessadé may be colonized with flies that are locally adapted to savannah landscapes, as opposed to tsetse flies from Coastal Guinea that are locally adapted to mangrove environements (De Meeûs et al., 2015). Local adaptation may considerably limit gene flow between these two lanscapes. Densities also seemed much smaller in the Guinean mangrove as compared to those found in the present work. Whatever the causating agent, dispersal and density can be negatively correlated in tsetse flies (De Meeûs et al., 2019). For the closely related *G. p. palpalis*, in forests of Cameroon or Côte d’Ivoire, dispersal appeared in the same order of magnitude (1-2 km/generation), but with much higher densities (50-200 flies / km²).

On the other hand, *G. m. submorsitans*, displayed a very scarce distribution across a rather important zone, and an absence of subdivision signature, suggesting a free dispersal at a very large scale (80 km) per generation (i.e. every two to five months). This may be due to the low power of the test due to the small subsample size observed in Keur-Tamba (seven individuals). Nevertheless, the only significant subdivision we found (*p*-value=0.0117) was between this area and Koutanga (43 individuals) (34 km), while Missirah (140 individuals) and Koutanga (42 km) gave a non-significant differentiation (*p*-value=0.3611). So, low power does not represent a plausible explanation, and even without Keur-Tamba, 42 km/generation between Missirah and Koutanga still remains a considerable dispersal distance. Effective population density appeared difficult to estimate given the absence of information on where this species is present. This is why our estimate varied from 1 to 8 tsetse flies per km² on average, and ranging from 0 to 16. Very large zones remained unexplored, in particular in Gambia. Given the dispersal capabilities of this species, and the variability of capture success from one spot to the other (See Table S1 and Figure S1), more needs to be done before any definitive conclusions can be inferred about the effective population density of this species in Southern Senegal and Gambia. Additionally, our observations were based on a rather limited number of loci, with rather substantial frequencies of null alleles, which may have contributed to some degree of uncertainty.

However, some published studies seemed to present smaller dispersal for *G. m. submorsitans*. A reinvasion study estimated the dispersal at 1 km/generation in Nigeria (Riordan, 1976). Population genetics studies were reassessed to try to avoid temporal issues, as far as we could. Details of these reanalyzes are presented in the Appendix D. In Gambia and Ethiopia (Krafsur et al., 2000; Krafsur & Endsley, 2002), this subspecies display long enough dispersal distances (around 20 km/generation), though apparently smaller than what we observed in Southwest Senegal. Extending to closely related subspecies, an MRR study of *G. m. morsitans* in Rhodesia found a much smaller possible dispersal of around 1 km/generation (Dame et al., 1975). Some other papers on the same subspecies (Krafsur & Endsley, 2002; Ouma et al., 2007) could not be used as explained more thoroughly in the appendix D. Nevertheless, a paper from flies from Zambia and Malawi (Nakamura et al., 2019), which we entirely reanalyzed (Appendix D), produced impressive dispersal distances in absence of physical barriers, with at least 95 km/generation and an absence of significant genetic differentiation at 571 km distance, while high montains seemed to restrict such dispersal to a still impressive 18 km/generation. Finally, results on *G. m. centralis* (Krafsur et al., 2001; Krafsur & Endsley, 2002) could not be used, as explained in Appendix D. It seems that, depending on the circumstances, the very large dispersal distances found in our paper, can, at least, be met in that species, but also in some other species, when no geographic barriers exist (Nakamura et al., 2019; Kagbadouno et al., 2024; Brito et al., 2025; De Meeûs et al., 2025), and particularly so during the rainy season (Cuisance et al., 1985).

While random mating could be safely assumed in *G. p. gambiensis* sub-populations, heterozygote deficits found in *G. m. submorsitans* suggested the existence of sib-mating. Sib-mating may result from a combined effect of huge geographic distances between larviposition sites (assuming those correspond to the ultra-dense pockets we found), systematic mating of imagoes within these sites just after the emergence, and large dispersal of some individuals. Nevertheless, the limited number of loci and relatively high levels of null allele frequencies bring some doubts on the real cause, and even the reality of this heterozygote deficit.

There was no signature of sex-biased dispersal in *G. p. gambiensis*. For *G. m. submorsitans*, further analyses with more subsamples outside the zone studied, in particular in Gambia, and more numerous and more performant markers will be required to investigate this issue.

## Conclusions

Despite the important population differences observed in these two species from Southern Senegal, similar recommendations may be produced from our results. The population of *G. p. gambiensis* appeared rather large, dense, and continuously structured into connected small subpopulations. Moreover, many potential sites remained unexplored, in particular in the Gambia. The population of *G. m. submorsitans* displayed spectacular capacities for dispersal (around 80 km / generation at least). Such dispersal capacities may appear prohibitively long as compared to what is currently admitted for tsetse flies (Hargrove, 2004). Nevertheless, some mark-release recapture studies suggest this may be possible, and particularly so during rainy seasons (Cuisance et al., 1985). Cuisance et al. (1985) indeed noted that most dispersive *G. p. gambiensis* of his experiment in Burkina Faso may travel 4.4 km/day (i.e. potentially 260 km during a life time if we count two months for one generation), with one fly traveling 22 km in only five days. Moreover, as for *G. p. gambiensis*, many potential spots may exist for *G. m. submorsitans* outside the zone we surveyed, particularly so in the Gambia, as observed in other surveys (Krafsur et al., 2000; Krafsur & Endsley, 2002). According to these results, eliminating the populations of these two species in Southern Senegal thus does not seem as a realistic option so far. The rolling carpet strategy used in the Niayes against *G. p. gambiensis* in Northern Senegal began in 2011 (Vreysen et al., 2021), and was considered successful in 2022 (Seck et al., 2024), i.e. after 11 years. The surface involved covered 1,000 km² for a cost estimated at 6.4 millions of euros in 2016 (i.e. an estimation fo 1,000 euros per year per km²) (Bouyer et al., 2014). This would be hard to apply for the populations of tsetse flies involved in the present study. Indeed, the surfaces targeted would probably outreach 4,000 km² and 10,000 km² for *G. p. gambiensis* and *G. m. submorsitans* respectively and expected costs between 4 and 10 billions of euros per year. Integrated, targeted and continuous pest management then appears to be the best option. For instance, continuous trapping has been shown to efficiently protect from trypanosome infections in Guinea, where vector control lightly (if any) affects the population of *G. p. gambiensis* as a whole (Courtin et al., 2015; Kagbadouno et al., 2024).

It appeared that classic microsatellite markers used in the present study showed some limitations, especially for *G. m. submorsitans*. Adapting new typing strategies, as SSRseq markers (Lepais et al., 2020), may allow getting more markers, with less amplification problems and all located in autosomes. These markers are currently being developed on several species of the palpalis group, and the first results are promising (De Meeûs et al., 2025).

Finally, the present results on population genetic structure of *G. p. gambiensis* and *G. m. submorsitans* in Southwest Senegal advocate that localized integrated pest management is likely to be the most effective approach for controlling tsetse and reducing trypanosome transmission. However, further entomological, parasitological, species-distribution modelling and socio-economic investigations and the use of more informative genetic markers are needed to obtain a more precise understanding of the epidemiological processes involved.

## Appendices

### Appendix A: When and which false discovery rate controlling procedure to use in repeated test series

When the *k* tests of a series are independent, the Benjamini and Hochberg FDR correction (Benjamini & Hochberg, 1995, 2000) (BH) applies. Under dependency, one would be inclined using Benjamini and Yekutieli procedure (Benjamini & Yekutieli, 2001) (BY). Nevertheless, BY is highly conservative, and maybe prohibitively so (Reiner et al., 2003), especially so for tests on proportions (De Meeûs, 2014). Moreover, for positively or moderately negatively dependent test statistics, BY presents no real FDR advantages as compared to BH, the use of which is then more suitable (Benjamini & Yekutieli, 2001; Reiner et al., 2003). Negatively dependent tests means that the strength of H1 (with a better chance of small *p*-values) of some tests in the series implies that other tests in the series will tend to have also small *p*-values, while other sets of tests are undertaken under H0. Typically, in a post-hoc multiple testing after an analysis of variance or after multiple average comparisons, we expect such moderate negatively correlated *p*-values. Nevertheless, in some instances, such correlation may be strong enough to violate the assumptions needed to apply BH. This would be the case, for population genetics analyses, when different sets of loci are in strong linkage disequilibrium (LD) and others are not. This is also what would be observed during differentiation tests between different pairs of subsamples taken from a population displaying isolation by distance, with a lot of pairs highly significant because from remote sites, and other pairs not so different (only by chance) because from close by sites. In such instances, detecting the true significant tests would require the use of BY. In all other cases, BH is expected to give more powerful, yet still reasonably robust, results. We can also notice here how more relevant a unique isolation by distance test would be as compared to the multiple testing of subdivision between paired subsamples.

### Appendix B: Scripts for geosphere measures

#### Computing geographic distances

LongLat1 <- read.table("LongLat1.txt", header=TRUE)
LongLat2 <- read.table("LongLat2.txt", header=TRUE)
distGeo(LongLat1,LongLat2)
TabDgeo<-data.frame(distGeo(LongLat1,LongLat2))
write.table(TabDgeo,"DistGeo.txt", col=NA, sep="\t", dec=".")

LongLat1.txt is a text file with GPS coordinates of the first subsample of each pair between which distance needs being measured. LongLat2.txt is a text file with GPS coordinates of the second subsample of each pair between which distance needs being measured. GPS coordinates are in decimal degrees in two columns in the order Longitude and Latitude.

The results are written in the file DistGeo.txt, in one column, in which each line is in the same order as in the two input files.

#### Computing surfaces

Dataset <- read.table("Traps1FlyT0-3LongLat.txt", header=TRUE)
areaPolygon(Dataset)

Traps1FlyT0-3LongLat.txt is a text file with the GPS coordinates in decimal degrees of each location used to define the targeted polygon. The file has two columns corresponding to the longitude and latitude of each location, in that order. The defined perimeter must be smooth without any loop. The package generates the surface in m².

### Appendix C: Scripts for sex-biased dispersal analyses

#### For G. p. gambiensis autosomes

GpgSineSaloumSexBias<-read.table("GpgSineSaloum6LociSexBiasData.txt",header=T) mean(AIc(GpgSineSaloumSexBias[,-2])[GpgSineSaloumSexBias[,2]=="F"]) mean(AIc(GpgSineSaloumSexBias[,-2])[GpgSineSaloumSexBias[,2]=="M"])
sexbias.test(dat = GpgSineSaloumSexBias[, −2], sex = GpgSineSaloumSexBias[, 2], nperm = 10000, test = "mAIc")
var(AIc(GpgSineSaloumSexBias[,-2])[GpgSineSaloumSexBias[,2]=="F"]) var(AIc(GpgSineSaloumSexBias[,-2])[GpgSineSaloumSexBias[,2]=="M"])
sexbias.test(dat = GpgSineSaloumSexBias[, −2], sex = GpgSineSaloumSexBias[, 2], nperm = 10000, test = "vAIc")
wc(GpgSineSaloumSexBias[GpgSineSaloumSexBias[,2]=="F",-2]) wc(GpgSineSaloumSexBias[GpgSineSaloumSexBias[,2]=="M",-2])
sexbias.test(dat = GpgSineSaloumSexBias[, −2], sex = GpgSineSaloumSexBias[, 2], nperm = 1000, test = "FST")
sexbias.test(dat = GpgSineSaloumSexBias[, −2], sex = GpgSineSaloumSexBias[, 2], nperm = 1000, test = "FIS")

#### For G. m. submorsitans

Gms5LociArea<-read.table("Gms5LociAreasSexBiasData.txt",header=T) mean(AIc(Gms5LociArea[,-2])[Gms5LociArea[,2]=="F"]) mean(AIc(Gms5LociArea[,-2])[Gms5LociArea[,2]=="M"])
sexbias.test(dat = Gms5LociArea[, −2], sex = Gms5LociArea[, 2], nperm = 10000, test = "mAIc") var(AIc(Gms5LociArea[,-2])[Gms5LociArea[,2]=="F"])
var(AIc(Gms5LociArea[,-2])[Gms5LociArea[,2]=="M"])
sexbias.test(dat = Gms5LociArea[, −2], sex = Gms5LociArea[, 2], nperm = 10000, test = "vAIc") wc(Gms5LociArea[Gms5LociArea[,2]=="F",-2])
wc(Gms5LociArea[Gms5LociArea[,2]=="M",-2])
sexbias.test(dat = Gms5LociArea[, −2], sex = Gms5LociArea[, 2], nperm = 1000, test = "FST") sexbias.test(dat = Gms5LociArea[, −2], sex = Gms5LociArea[, 2], nperm = 1000, test = "FIS")

### Appendix D: Reanalyzes of population genetics data of published papers on *Glossina morsitans* ssp

For all papers, we undertook Rousset’s regression in order to extract the neighborhood size (*Nb*) from the reverse of the slope. We computed geographic distances (*D*_geo_) from GPS coordinates with the R-package geosphere and used *F*_ST_ measured between paired subsamples as given in the cited papers. Please note that some of the GPS coordinates were erroneous and we corrected those using GoogleEarth Pro. We then considered *Nb* as a proxy for the upper limit of the effective subpopulation size (*N_e_*). We used the *F*_ST_ measured between each subsample pair to compute the number of immigrants coming from each subsamples *N_e_m* and *m* the immigration rate as *m*=*N_e_m*/*N_e_*. Maximum value for *m* was 1, and null or negative *F*_ST_ were assigned *m*=1 also. Dispersal was then roughly inferred as *δ*=*mD*_geo_. Please note that whenever possible, we excluded subsample pairs that were not contemporaneous, that the precise date of sampling at two months precision (generation time of tsetse flies) was never available, and that *F*_ST_ measures could never be corrected for null alleles. The results are thus, at best, only indicative as orders of magnitude.

#### Glossina morsitans submorsitans

The first dataset was from Krafsur et al. (2000). Data from Gambia were from year 1997, containing potentially six cohorts, and from Ethiopia (late 1997). Details on sampled sites are presented in the Table AD1.

**Table AD1.**
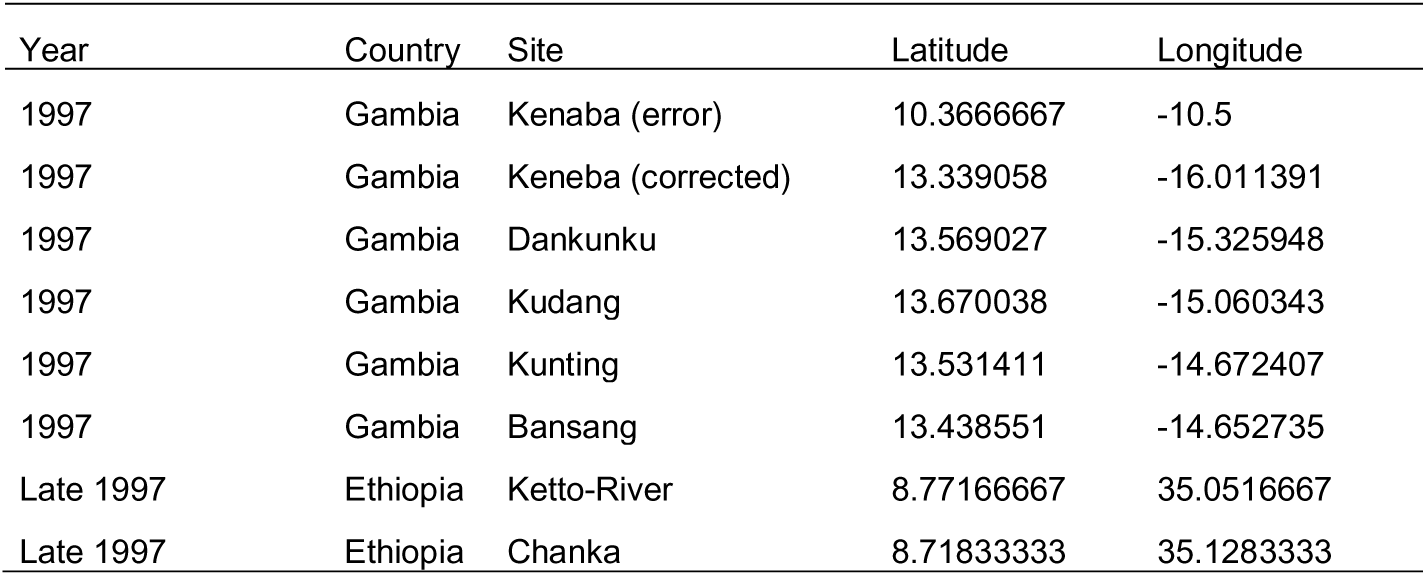
Date of sampling (year) Country and site sampled and GPS coordinates of *Glossina morsitans submorsitans* from Gambia and Ethiopia (Krafsur et al., 2000). Kenaba coordinates corrected with Google Earth Pro.

| Year | Country | Site | Latitude | Longitude |
| --- | --- | --- | --- | --- |
| 1997 | Gambia | Kenaba (error) | 10.3666667 | -10.5 |
| 1997 | Gambia | Keneba (corrected) | 13.339058 | -16.011391 |
| 1997 | Gambia | Dankunku | 13.569027 | -15.325948 |
| 1997 | Gambia | Kudang | 13.670038 | -15.060343 |
| 1997 | Gambia | Kunting | 13.531411 | -14.672407 |
| 1997 | Gambia | Bansang | 13.438551 | -14.652735 |
| Late 1997 | Ethiopia | Ketto-River | 8.77166667 | 35.0516667 |
| Late 1997 | Ethiopia | Chanka | 8.71833333 | 35.1283333 |

Subdivision measures obtained from mitochondrial (mt) between "contemporaneous" subsample pairs provided a rather good model (Figure AD1) with a slope =0.0167. It can be seen that including pairs between non-contemporaneous subsamples (between 1997 and late 1997 subsamples) would produce an inappropriate result. Alternatively, "contemporaneous" subsamples produced a regular slope, which suggest that 1997 subsamples on one hand, and late 1997 subsamples on the other hand came from more or less the same cohort. This slope led to a neighbourhood *Nb*=60 (and assumed *N_e_* of an order of magnitude of *N_e_*=50). Computing dispersal distances as indicated for each "contemporaneous" subsample pairs lead to an average *δ*=24 km/generation.

**Figure AD1.**
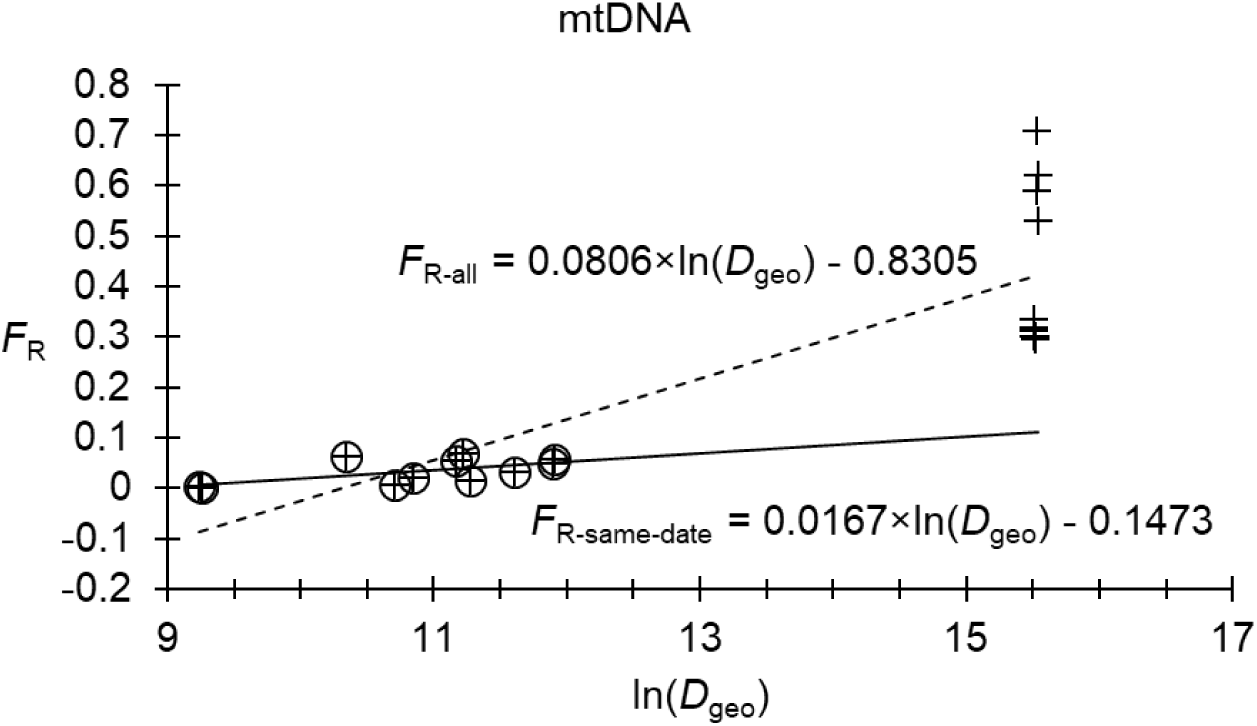
– Rousset’s model for isolation by distance for *Glossina morsitans submorsitans* between 1997 and between late 1997 subsamples (*F*_R-same-date_: more or less contemporaneous) (empty circles and plain line) and between all subsample pairs (crosses, dotted line).

The second dataset concerned the same Gambian subsamples but with microsatellite markers (Krafsur & Endsley, 2002). We obtained comparable results (Figure AD2), with *Nb*=50 (*N_e_*≈40) and *δ*=16 km/generation.

**Figure AD2.**
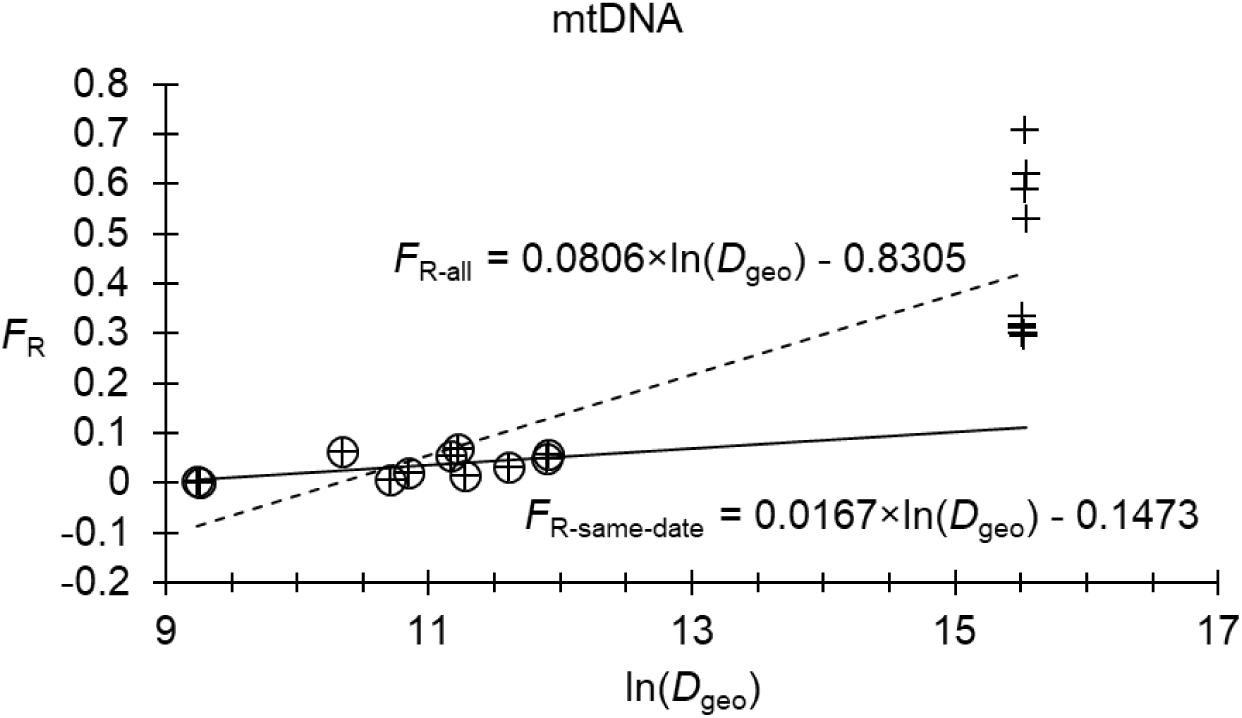
Rousset’s model for isolation by distance for *Glossina morsitans submorsitans* between 1997 and between late 1997 subsamples (*F*_R-same-date_: more or less contemporaneous) (empty circles and plain line) and between all subsample pairs (crosses, dotted line).

These figures were compatible with low densities: 0.01 and 0.02 flies/km² for microsatellite loci and mtDNA respectively, with a surface between 3,000 km² (microsatellite markers) and 6,000 km² (mtDNA). For the record, the surface defined by the sites is about 2300 km², and the one defined by these sites and those with *G. m. submorsitans* in Senegal reported here is around 4200 km². Nevertheless, the impossibility to correct for null alleles and the ignorance of the precise dates of sampling makes those results questionable and we may retain that dispersal distances appear quite important in that subspecies, in the Senegal-Gambian ecosystems.

#### Glossina morsitans morsitans

A study of mtDNA and microsatellites loci was undertaken from samples from Zambia, Mozambique and Zimbabwe along years 1995 and 1996 (i.e. across 12 generations of tsetse flies) (Krafsur & Endsley, 2002; Ouma et al., 2007) and four sites in Tanzania during 2004 (6 generations) (Ouma et al., 2007). Nevertheless, in addition to dates of sampling, some GPS coordinates appeared inaccurate, leading to discordance between GPS coordinates and geographical distances reported in the text. For instance, sites Lumbe and Usinga are said to be 12 km apart, but their GPS coordinates locate those at 21 km. Or, for Kilimamzinga and Rekomitjie (917 km in the text), woud be 1476 km aparts according to GPS. This is probably why we obtained unusable isolation by distance regressions.

In another and more recent study (Nakamura et al., 2019), raw data were made available and we reanalyzed entirely these data. Tsetse flies were sampled at three different dates corresponding to 29 generations: in Zambia in 2012 in the Lower Zambezi National Park (LZNP), in 2017 in the Musalungu Game Management Area (MGMA) and in Shikabeta; in Malawi in 2018 in the Nkhotakota Wild Life Reserve (NWR) and the Kasungu National Park (KNP). These corresponded to cohorts C0, C26 and C28 respectively.

The *F*_IS_ and *F*_ST_ are presented in the Figure AD3. Significant heterozygote deficits were well explained by null alleles, as the regression *F*_IS_∼*p*_nulls_ outputted a *R*²=0.96 and an intercept *F*_IS-0_=-0.063 in 95%CI=[-0.248, 0.1009]. Nevertheless, four loci needed to be removed from the analyses (Figure AD3). Locus Gmm15, was not well explained by null alleles, and displayed spatial and/or temporal disruptive selection, with allele 196 (blue in Figure AD3) almost fixed in NWR_C28, while absent elsewhere. Locus GmmL11 also displayed such a disruptive selection. Locus GmmP07 was not well explained by null alleles, stuttering or SAD. Finally, GmmL17 also displayed spatial and/or temporal disruptive selection, with allele 302 (yellow in Figure AD3) absent everywhere but in NWR_C28 with frequency 0.6. Note that locus GmmA06 displayed significant stuttering and was corrected with the pooling of alleles 175 to 193 into allele 173 and 197 to 214 into allele 195.

Please, note that problems probably arose during the analyses of Nakamura et al as they reported very important excesses of heterozygotes, which are not compatible with the dataset published in dryad that we used. The following results were thus obtained with 6 loci: GpCAG1, GpC101, GpC10b, GmmK22, GmmA06 (corrected for stuttering), and GmmC17.

**Figure AD3.**
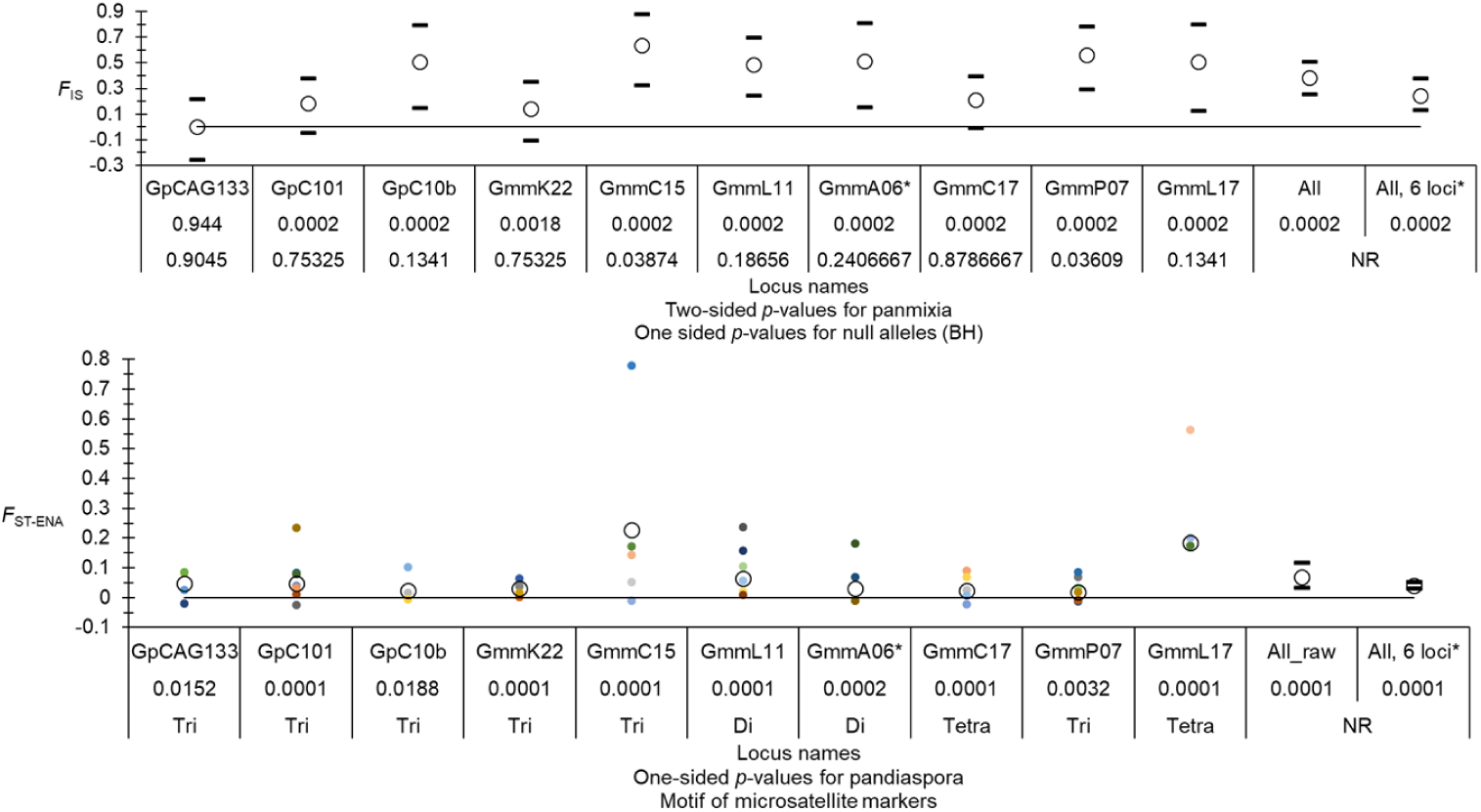
– Values of *F*_IS_ (top graphic) and *F*_ST -ENA_ (corrected for null alleles) (bottom), observed after the reanalysis of *Glossina morsitans morsitans* populations of Zambia and Malawi (Nakamura et al., 2019). Averages (empty circles) are given for each locus and over all loci (All). The 95% confidence intervals are presented by black dashes. Alleles with frequencies above 0.05 are also represented with their Weir and Cockerham’s estimator of *F*_ST_.

Subdivision, corrected for null alleles, could be measured between MGMA and Shikabeta (cohort C26), and between NWR and KNP (cohort C28). There was a non-significant subdivision between MGMA and Shikabeta with *F*_ST-ENA_=0.0017 in 95%CI=[-0.0079, 0.0107] (*p*-value=0.1814), which are 571 km distant from each other’s, as the crow flies, and a highly significant one between NWR and KNP, with *F*_ST-ENA_=0.0508 in 95%CI=[0.0194, 0.008780] (*p*-value<0.0001), which are 109 km distant. As mentioned in the article, NWR is isolated from other sites, and in particular from KNP, by mountains that probably represent efficient barriers. Average effective population sizes were *N_e_*=15 in minimax=[2, 139] for C26 subsamples, and *N_e_*=28 in minimax=[2, 283] for C28 subsamples. These figures lead to an average dispersal distance of 571 km / generation according to the first result, with a minimum of 95 km/generation. For the second pair, separated by mountains, dispersal was 18 km/generation on average, in 95%CI=[10, 49] and a minimax=[1, 109].

#### Glossina morsitans centralis

Six sites were sampled and analyzed with mt DNA and microsatellite loci in Zambia (two sites), Namibia, (one site), and Botswana at unknown dates of sampling and hard to locate sites (Krafsur et al., 2001; Krafsur & Endsley, 2002). This led to the impossibility to estimate neighborhood sizes as isolation by distance slopes were unreliable (close to 0), as a probable consequence of uncorrected null alleles, unreliable site locations, and probable prohibitive temporal distances between the different sites. On average, mtDNA and microsatellite loci outputted *F*_ST_=0.5861 and *F*_ST_=0.14 respectively, for "geographic" distances of 463-469 km on average. This is far above what we would have been expected with our data on *G. m. submorsitans* from Southern Senegal, but is the same order of magnitude of spatio-temporal subdivision found for *G. m. submorsitans* by the same authors (Figures AD1 and AD2). It is also four times what we found for spatio-temporal differentiation found for *G. m. morsitans* (Nakamura et al., 2019), with *F*_ST_=0.0318 and average distances of 391 km.

## Acknowledgements

Genotyping was undertaken in the MGX platform of the Cirad of Montpellier (UMR AGAP).

## Author contributions

MTB conceived the study, did the sampling, genotyping and corrected the manuscript. AGF participated to the sampling and corrected the manuscript. AS participated to the conception of the study, to genotyping and corrected the manuscript. MTS participated to the sampling and corrected the manuscript. AST participated to the sampling, to genotyping and corrected the manuscript.GG participated to the conception of the study and corrected the manuscript. SR participated to the conception of the study, to genotyping, corrected the manuscript and supervised the project.TdM participated to the conception of the study, to data analyses wrote the manuscript and supervised the project.

## Data, scripts, code, and supplementary information availability

Scripts of R are given in Appendices A and B. All data are available in Supplementary Files S1, S2 and S3, and in Supplementary Figure S1

---

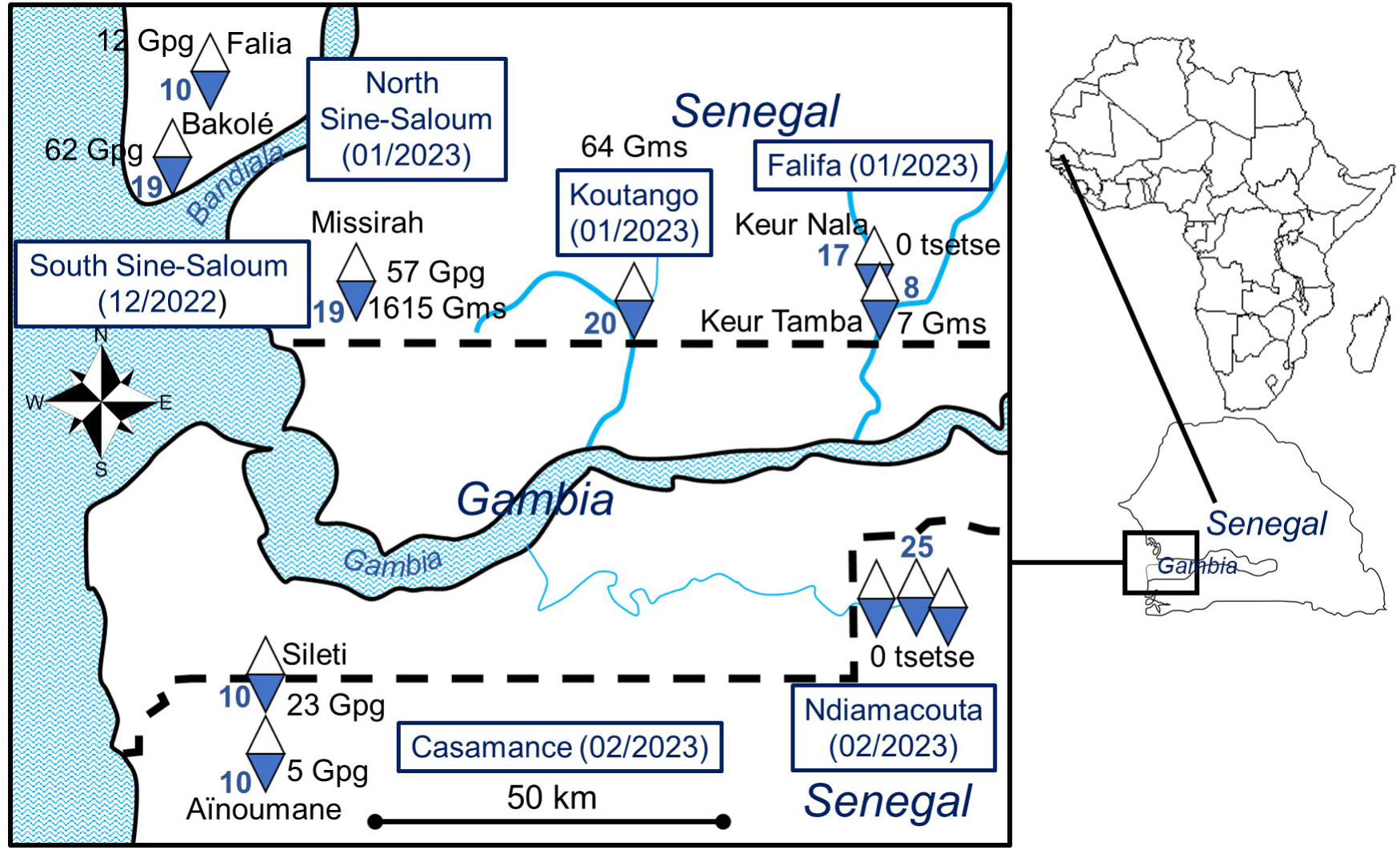

